# Immunomodulatory effects of tumor Lactate Dehydrogenase C (LDHC) in breast cancer

**DOI:** 10.1101/2024.11.26.625341

**Authors:** Adviti Naik, Remy Thomas, Aljazi Al-Khalifa, Hanan Qasem, Julie Decock

**Author notes:** Corresponding author: <u>Dr Julie Decock,</u> Qatar Biomedical Research Institute (QBRI), HBKU, Research and Development complex, Education City, Qatar Foundation, PO 34110, Doha, Qatar.

## Abstract

**Background:** Immunotherapy has significantly improved outcomes for cancer patients; however, its clinical benefits vary among patients and its effectiveness across breast cancer subtypes remains uncertain. To enhance its efficacy, it is important to gain more insight into tumor-intrinsic immunomodulatory factors that could be used as therapeutic targets. We previously identified Lactate Dehydrogenase C (LDHC) to be a promising anti-cancer target due to its role in regulating cancer cell genomic integrity. In this study, we investigated the effects of tumor LDHC expression on immune responses.

**Methods:** TIMER AND TIDE deconvolution methods were used to investigate the relationship between tumor *LDHC* expression, immune cell infiltration and T cell dysfunction. Multiplex cytokine assays and flow cytometry analyses of breast cancer cell monocultures, and direct and indirect cancer cell-immune cell co-culture models were performed to assess the effect of LDHC knockdown on the secretion of inflammatory mediators and the expression of immune checkpoint molecules. T cell activity was determined by IFN-γ ELISPot assays and 7-AAD viability flow cytometry of cancer cells in direct co-culture.

**Results:** TIMER and TIDE analyses revealed that tumor *LDHC* expression is associated with T cell dysfunction in breast cancer and worse post-immunotherapy survival in melanoma. Depletion of LDHC in three breast cancer cell lines (MDA-MB-468, BT-549, HCC-1954) enhanced T cell activation and cytolytic function (4-hour direct co-culture). Analysis of cancer cell monocultures revealed an increase in secreted pro-inflammatory cytokines (IFN-γ, GM-CSF, MCP-1, CXCL1), a decrease in immunosuppressive factors (IL-6, Gal-9) and a reduction in tumor cell surface PD-L1 expression following LDHC knockdown. Using 72-hour direct co-cultures with LDHC-silenced cancer cells, we observed a decrease in tumor-promoting cytokines (IL-1β, IL-4 and IL-6) and an increase in the tumor-inhibiting cytokine CXCL1. Furthermore, LDHC knockdown reduced the number of CD8+ T cells expressing PD-1 and CTLA-4, as well as the cell surface expression of CTLA-4, TIGIT, TIM3, and VISTA.

**Conclusions:** Our findings suggest that targeting LDHC may improve anti-tumor immune responses by modulating the secretion of pro- and anti-tumorigenic cytokines and impairing immune checkpoint signaling. Further studies are needed to elucidate the molecular mechanisms by which LDHC modulates these responses in breast cancer.

## BACKGROUND

Tumor development and progression is a multifactorial process that involves genetic, environmental and lifestyle factors that drive cellular transformation, shape the tumor microenvironment and impact the anti-tumor immune response. Under normal physiological conditions, the innate and adaptive immune responses coordinate to detect and eliminate threats by pathogens and abnormal or transformed cells. The ability of tumor cells to escape immune recognition and killing is known as the escape phase of the immunoediting concept, which is preceded by the equilibrium phase, during which escaping tumors evolve to express and secrete various factors that promote an immunosuppressive milieu [1]. Dysregulation of the anti-tumor immune response has been directly correlated with tumor progression and poor prognosis in many solid tumors, spurring the development of immunotherapy as a therapeutic approach to reinvigorate and reprogram the host immune response [2]. In particular, immune checkpoint blockade has shown significant clinical benefit in specific cancer types such as melanoma and non-small cell lung cancer [3,4]. To date, eight immune checkpoint inhibitors targeting CTLA- 4 (ipilimumab), PD-1 (pembrolizumab, nivolumab, cemiplimab, dostarlimab), and PD-L1 (atezolizumab, durvalumab, avelumab) have received FDA approval for the treatment of unresectable or metastatic solid tumors, including triple negative breast tumors [5,6]. Despite their ability to induce durable responses, the majority of patients do not exhibit clinical benefit from immune checkpoint blockade due to various immune evasion mechanisms [7]. Therefore, combinatorial strategies are needed to improve clinical response. In this context, the identification and targeting of tumor-intrinsic oncogenic mediators that promote immunosuppression is of great strategic interest.

Cancer testis antigens (CTAs) are a group of proteins with testis-restricted (expressed only in testis) or testis-selective (expressed in testis and up to two other tissues) expression in normal tissues and aberrant expression in numerous cancers [8,9]. The highly tumor restricted expression of CTAs, along with their immunogenic properties and multifaceted roles in oncogenic processes positions CTAs as excellent candidates for therapeutic targeting. Lactate dehydrogenase C (LDHC, LDHX, CT32) is a CTA with aberrant expression in several solid tumors, including breast, lung, renal and colon cancer, which exhibits immunogenic properties [8,10]. It is a member of the lactate dehydrogenase (LDH) family, which catalyzes the interconversion of pyruvate and lactate. While LDHA and LDHB form heterotetramers with ubiquitous expression in skeletal muscle and the heart, LDHC exists as a homotetramer with testis-restricted expression, playing a critical role in regulating sperm energy metabolism and motility [11,12]. In tumor cells, LDHC expression is regulated by the Sp1 and CREB transcription factors, in addition to promoter CpG island hypomethylation [13]. LDHC levels in serum and serum-derived exosomes from patients with breast and hepatocellular carcinoma positively correlate with cancer diagnosis, tumor size, and recurrence [14,15]. Furthermore, *LDHC* expression in renal tumor tissue is associated with shorter survival [16]. In line with its function in the testis, LDHC is a key regulator of aerobic glycolysis and has also been shown to promote tumor cell invasion, migration, and proliferation through activation of the PI3K/Akt/GSK-3B signaling pathway [17,18]. We recently demonstrated that silencing of LDHC in breast cancer cells leads to DNA damage accumulation and dysregulation of the cell cycle, impairing long-term cancer cell survival [19]. Hence, LDHC can be included alongside PBK, SSX2 and MAGE-C2 as one of the 27 genomic integrity-regulating CTAs [20].

In this study, we explored whether LDHC expression in tumor cells may affect the tumor microenvironment, particularly the intricate dynamic interplay between tumor cells and immune cells that drive immune escape. Previously, we observed an increased expression of *LDHC* in basal-like and Her2-enriched breast tumors compared to the less aggressive luminal and normal-like subtypes [19], suggesting that LDHC may play a more prominent role in basal-like or Her2-enriched tumors, which are associated with more immunogenic tumor microenvironments [21]. To further explore this, we examined whether LDHC expression in basal-like and Her2-enriched breast cancer cells impacts immune cell function. Utilizing co-culture models, we found that LDHC expression in breast cancer cells may promote an immunosuppressive tumor microenvironment and immune cell dysfunction by altering the secretion of cytokines and regulating the expression of various immune checkpoints. Similarly, we demonstrated that aberrant expression of the cancer testis antigen PReferentially expressed Antigen in Melanoma (PRAME) in tumor cells contributes to dampening of the adaptive immune response, suggesting that at least two CTAs, LDHC and PRAME, may impact tumor development and progression through modulation of anti-tumor immunity [22]. These findings indicate that cancer testis antigens could serve as promising therapeutic targets, potentially impairing pro-tumorigenic intrinsic mechanisms and mitigating an immunosuppressive microenvironment. As such, our results advocate for the development of combinatorial therapies to target CTAs in combination with immune-based interventions.

## METHODS

### Cell culture

MDA-MB-468, HCC-1954 and BT-549 cell lines were purchased from the American Tissue Culture Collection (ATCC). MDA-MB-468 cells were maintained in Dulbecco’s Modified Eagle’s Medium (Gibco-BRL) supplemented with 10% (v/v) Fetal Bovine Serum (FBS) (Hyclone US origin, GE Lifescience), 50 U/ml penicillin and 50 μg/ml streptomycin (Gibco-BRL). HCC-1954 cells were maintained in ATCC-formulated Roswell Park Memorial Institute (RPMI) 1640 medium (Gibco-BRL) supplemented with 10% (v/v) FBS, 50 U/ml penicillin and 50 μg/ml streptomycin. BT-549 cells were maintained in ATCC-formulated RPMI 1640 medium supplemented with 10% (v/v) FBS, 50 U/ml penicillin and 50 μg/ml streptomycin (Gibco-BRL), and 0.023 IU/ml insulin (Sigma-Aldrich). All cell lines were maintained in a humidified incubator at 37 °C, 5% CO2 and regular mycoplasma testing was performed using a PCR-based assay.

### LDHC silencing

LDHC siRNA #1–4 smartpool (siGENOME SMARTpool) and control siRNA #1–4 smartpool (siGENOME non-targeting siRNA Pool#1, D-001206-13-20) were purchased from Dharmacon. Adherent cancer cells were transfected at 60–70% confluency with 20 nM siRNA pools and Lipofectamine RNAiMAX (Thermo Fisher Scientific) following the manufacturer’s instructions. Knockdown of LDHC expression was assessed by real time qRT-PCR and western blotting.

### RNA extraction and quality assessment

Total RNA was isolated using the PureLink RNA Mini kit (Ambion) following the manufacturer’s protocol. The RNA quantity and purity were assessed by Nanodrop measurement. Reverse transcription of 1 µg RNA was performed using MMLV-Superscript reverse transcriptase (Thermo Fisher Scientific) and random hexamers (Applied Biosystems) resulting in a final concentration of 50 ng/µl cDNA.

### Quantitative real-time reverse transcription polymerase chain reaction

*LDHC* expression was quantified using specific 5′FAM-3′MGB Taqman gene expression primer/probe sets (Hs00255650_m1, Applied Biosystems). *PD-L1*, *CD80*, *GAL-9* and *PVR* expression was quantified by SYBR Green-based qPCR using PowerUp SYBR Green mastermix (Applied Biosystems) and primers designed through PrimerBLAST (NCBI). qRT-PCR was performed on the QuantStudio 7 system (Applied Biosystems). Relative expression levels were normalized to the housekeeping gene RPLPO using a Taqman gene expression primer/probe set (4333761F, Applied Biosystems) or primers designed for SYBR-Green based PCR (Forward primer sequence 5’-TCCTCGTGGAAGTGACATCG-3’, Reverse primer sequence 5-TGGATGATCTTAAGGAAGTAGTTGG-3’).

### Western blotting

Cell protein lysate was isolated using RIPA buffer (Pierce) supplemented with HALT protease and phosphatase inhibitor cocktail (Thermo Fisher Scientific). Western blotting was performed using a standard protocol as previously described [23]. Primary antibodies utilized include antibodies against LDHC (#ab52747, Abcam, 1:500), PD-L1 (#13684, Cell Signaling Technologies, 1:1000) and β-actin (#4970, Cell Signaling Technologies, 1:1000). Horseradish Peroxidase (HRP)-linked anti-rabbit/mouse secondary antibody incubation followed by enhanced chemiluminescent substrate (ECL) Supersignal West Femto (Pierce) incubation was used to visualize the protein bands of interest on the ChemiDoc XRS+ Imaging system (Biorad).

### Peripheral blood lymphocyte (PBL) isolation

Buffy coat samples, collected from healthy donors at the Hamad Medical Corporation Blood Donation Center, were diluted with Dulbecco’s Phosphate-Buffered Saline (DPBS, Gibco-BRL), layered on Lymphoprep™ (Stem Cell Technologies) and subjected to density gradient centrifugation to isolate the peripheral blood mononuclear cells (PBMCs). The PBMCs were washed and frozen in liquid nitrogen in freezing media (50% FBS, 40% serum-free Roswell Park Memorial Institute 1640 medium (RPMI), 10% Dimethyl sulfoxide) until further use. Prior to assays, PBMCs were defrosted in RPMI medium supplemented with 10% FBS and incubated overnight at 37°C and 5% CO2. To isolate the non-adherent peripheral blood lymphocyte population, the cells were plated in a flat-bottom multi-well plate (Thermo Fisher Scientific, Nunclon Δ Surface) and incubated for 2 h at 37°C and 5% CO2. Next, the non-adherent PBLs were activated overnight using 2 μg/ml of plate-bound anti-human CD3 and CD28 antibodies (eBioscience) at 37°C and 5% CO2. HLA typing of PBMCs was obtained for nine different HLA loci (A, B, C, DRB1, DRB3/4/5, DQA1, DQB1, DPA1, and DPB1) with one or two field resolution and donors with matched loci to MDA-MB-468, BT-549 and HCC- 1954 cancer cells were selected for further analysis.

### Indirect co-culture

A total of 5 × 10^4^ cancer cells were seeded per well in a 24-well plate. Next, activated PBLs were placed on top using transwell inserts with 0.4 µm pore size (Corning) and a Target:Effector (T:E) ratio of 1:20 to enable exchange of soluble factors between cancer cells and PBLs without direct cell-cell contact. Wells with PBLs alone were used as a negative control. The cells were co-cultured for 72 h at 37°C and 5% CO2, after which the PBLs, cancer cells and conditioned media were collected for flow cytometry and cytokine multiplex analysis.

### Direct co-culture

Activated PBLs were seeded at 1 × 10^6^ cells per well in a U-bottom 96-well plate. Cancer cells were labelled with 10 nM Qtracker™−655 (Thermo Fisher Scientific) according to manufacturer’s instructions. A total of 2x10^4^ cancer cells were added to the wells with activated PBLs (T:E ratio 1:50) and co-cultured at 37°C, 5% CO2. After 4 h of direct co-culture, the cells were harvested for interferon (IFN)-γ ELISpot assay and cytotoxicity analysis by flow cytometry. In addition, 72 h direct co-cultures of cancer cells and PBLs at T:E ratio of 1:20 were performed, and cells and conditioned media were collected for immune checkpoint flow cytometry and cytokine multiplex analysis.

### IFN-γ ELISpot

The Human IFN-γ ELISpot PLUS kit (HRP, Mabtech) was utilized according to manufacturer’s instructions. A total of 5 × 10^4^ PBLs from short-term direct (4 h) co-culture experiments were seeded per well. PBLs activated with 2 μg/ml of plate-bound anti-human CD3 and CD28 antibodies (eBioscience) or PBS served as positive and negative controls respectively. The wells were incubated at 37°C, 5% CO2 for 24 h and the number of IFN-γ spot forming units were quantified using an ELISpot reader (Autoimmun Diagnostika GmbH).

### Cytotoxicity analysis

After 4hr direct co-culture, cells were washed and resuspended in PBS, followed by staining with 7-Aminoactinomycin D (7-AAD) (eBiocience) at room temperature for 5 min. Flow cytometry was performed by recording 50,000 events/sample using the LSRFortessa X-20 instrument and FlowJo V10.10.0 software (BD Biosciences). Non-viable cancer cells were gated as positive for Qtracker and 7-AAD staining.

### Cytokine multiplex analysis

Cell culture supernatants from 72hr direct co-cultures were diluted 1:2 and used to determine the expression of 23 soluble proteins with the Bio-Plex Pro Human Cytokine 17-plex Assay (Biorad #M5000031YV) and a custom 6-plex Luminex array (R&D Systems, **Table S1**) according to manufacturer’s instructions using the Bioplex-200 system (Biorad). A 13-point standard curve was generated to extrapolate the soluble protein levels and the data was analyzed using the Bioplex Manager Software (Biorad).

### Flow cytometric analysis of immune checkpoint expression

PBLs and cancer cells from direct and indirect co-cultures were used for multi-marker flow cytometry to detect the expression of immune checkpoint markers. Cells were washed and resuspended in 100 μl of staining buffer containing Human Fc Block™ (564219, BD Biosciences). The expression of immune checkpoint receptors was determined using the following antibodies: PD-1 PE-Dazzle 594 (329940, Biolegend), VISTA BV421 (566750, BD Biosciences), CTLA-4 BV786 (563931, BD Biosciences), TIM3 BV650 (565565, BD Biosciences), LAG-3 PE (565617, BD Biosciences), TIGIT BUV395 (747845, BD Biosciences) and CD8 BV510 (563919, BD Biosciences). In parallel, we determined the expression of respective ligands: CD80 BV510 (740150, BD Biosciences), CD86 Alexa700 (564544, BD Biosciences), PD-L1 PE-Cy7 (558017, BD Biosciences), PD-L2 BV786 (563843, BD Biosciences), VISTA BV421 (566750, BD Biosciences), MHC-II BV650 (564231, BD Biosciences), GAL-9 PE (565890, BD Biosciences) and PVR BUV395 (748272, BD Biosciences). Flow cytometry was performed by recording 50,000 events/sample using the LSRFortessa X-20 instrument and FlowJo V10.10.0 software (BD Biosciences).

### Tumor immune infiltration, dysfunction and exclusion analyses

The Tumor IMmune Estimation Resource (TIMER) algorithm [24] was applied to investigate the relationship between tumor *LDHC* expression and immune cell infiltration (http://timer.cistrome.org/). In addition, we used the TIMER 2.0 platform to apply the Estimating the Proportions of Immune and Cancer cells (EPIC) algorithm which generates direct estimates of cell proportions by fitting single-cell RNA expression data to bulk RNAseq data. Finally, we utilized the Tumor Immune Dysfunction and Exclusion (TIDE) algorithm [25,26] to predict the effect of tumor *LDHC* expression on T cell dysfunction and immunotherapy response (http://tide.dfci.harvard.edu).

### Protein-protein interaction network and gene ontology analysis

Protein-protein interaction network analysis was performed using STRING online tool (http://www.string-db.org), and visualization was achieved using full STRING network with high confidence network edges. Gene ontology (GO) analysis was conducted using the Enrichr tool [27–29] and GO terms, involving multiple genes, were visualized using SRplot [30].

### Statistical analyses

Statistical analyses were performed using the Student’s t-test with p values ≤0.05 deemed being statistically significant. Data of at least three independent biological replicates are represented as mean ± standard error of mean (SEM), unless stated otherwise. Statistical analyses and data visualization were performed using GraphPad Prism v10.0.0.

## RESULTS

### Aberrant tumor *LDHC* expression is associated with T cell dysfunction

We investigated the relationship between aberrant *LDHC* tumor expression, immune cell infiltration, and T cell function in breast cancer using used various deconvolution methods. To gain more insight into potential associations of tumor *LDHC* expression with immune cell infiltration, we applied the Tumor IMmune Estimation Resource (TIMER) algorithm to estimate the relative abundance of six major immune cell types (B cells, CD8+ T cells, CD4+ T cells, macrophages, neutrophils and dendritic cells) in the TCGA breast cancer dataset **(Fig 1A)**. We observed a positive correlation between *LDHC* expression and B cell infiltration in Her2-enriched tumors, macrophages and dendritic cells in luminal B tumors, and CD4+ T cells in breast tumors at large. Next, we utilized the Estimating the Proportions of Immune and Cancer cells (EPIC) algorithm to account for tumor purity variety across samples, generating a more accurate direct estimate of cell proportions within tumors. Using EPIC, we did not find significant associations between *LDHC* expression and the infiltration of the six major immune cell subtypes previously analyzed using TIMER. However, we found a negative correlation between *LDHC* expression and NK cell infiltration in basal-like breast tumors, a cell type which is not available in the TIMER deconvolution method. Considering both methods, *LDHC* expression was found to be negatively associated with NK cell infiltration in basal-like breast tumors and positively associated with B cell infiltration in Her2-enriched breast tumors. In addition, both methods indicate that *LDHC* expression is positively correlated with tumor purity, suggesting that tumor cells are likely the main source of LDHC in the tumor microenvironment **(Fig 1A - bottom)**. To study whether LDHC expression affects immune function and thus clinical outcomes and responses to immunotherapy, we used the Tumor Immune Dysfunction and Exclusion (TIDE) algorithm. TIDE classifies tumors with high or low *LDHC* expression into two groups based on cytotoxic T lymphocytes (CTLs) abundance. While high CTL infiltration is commonly associated with good prognosis in cancer, we observed that high expression of *LDHC* in Her2-enriched and triple negative breast tumors negates or drastically reduces the favorable connotation of CTL infiltration with regards to overall survival, suggestive of T cell dysfunction which may in turn negatively impact immunotherapy responses **(Fig 1B)**. Due to the lack of available data on immunotherapy responses in breast cancer within the TIDE database, we explored the effect of *LDHC* expression on treatment response in patients with melanoma who received PD-1 immune checkpoint blockade. Similar to our observations in Her2-enriched and triple negative breast cancer, high expression of *LDHC* in melanoma significantly diminished the favorable association of CTL infiltration with overall survival **(Fig 1C)**. Furthermore, melanoma patients with high *LDHC* tumor expression exhibited a shorter overall and relapse-free survival in response to PD-1 blockade **(Fig 1D)**. Together, these findings suggest that while aberrant tumor expression of *LDHC* may not strongly affect immune cell infiltration, it is linked to T cell dysfunction.

**Figure 1.**
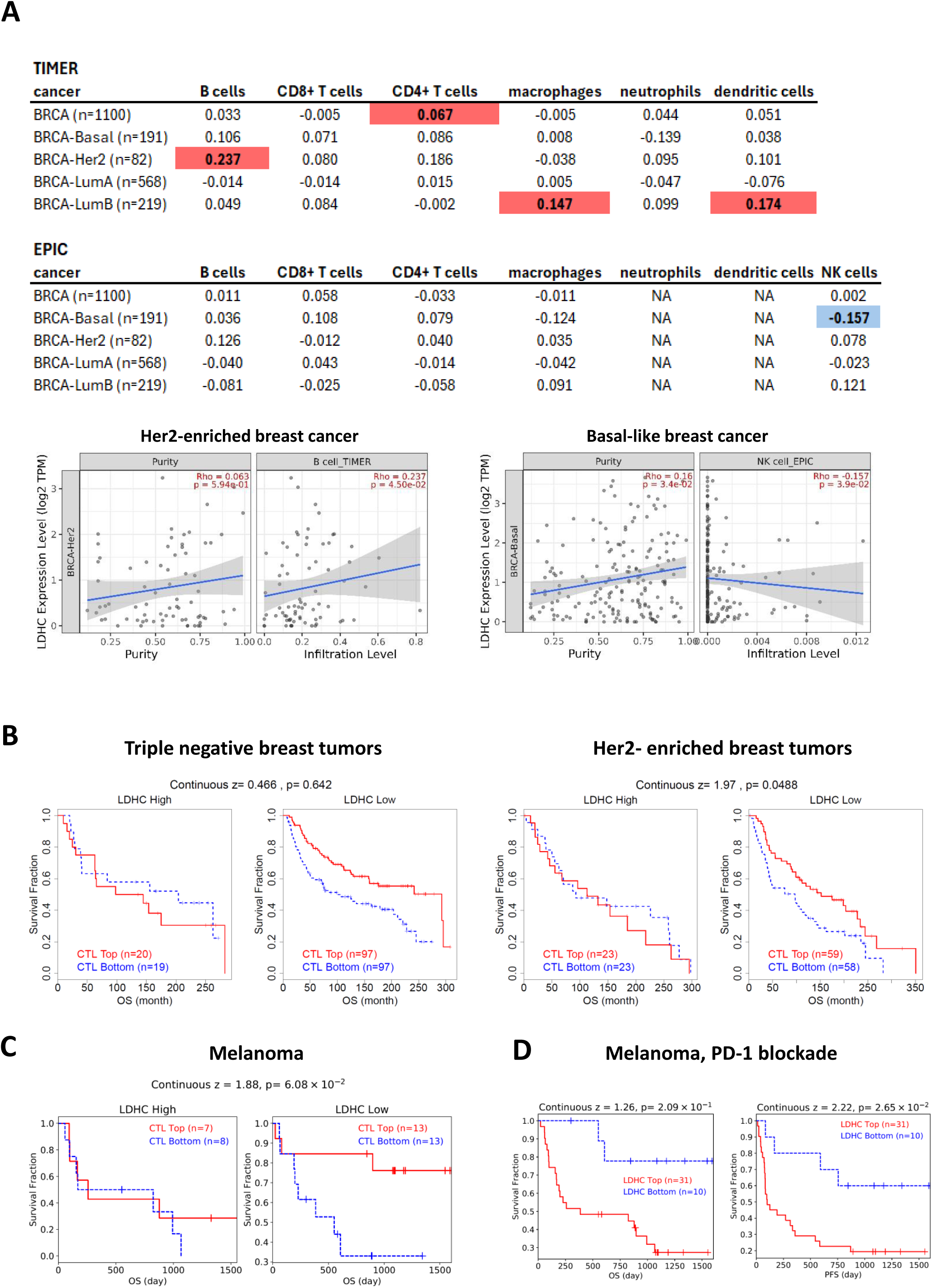
Association of tumor *LDHC* expression with immune cell infiltration and T cell dysfunction. **A)** Correlation of tumor LDHC expression and infiltration of immune cells subsets according to TIMER and EPIC deconvolution methods. Significant positive and negative spearman’s rho values are highlighted in red and blue respectively. Significant correlations in Her2-enriched and basal-like breast cancer are depicted in scatter plots. **B)** Kaplan Meier plots of triple negative and Her2-enriched breast cancer patients classified as having *LDHC* high or low expressing tumors and dichotomized by cytotoxic T lymphocyte (CTL) infiltration level. **C)** Survival analysis of melanoma patients with *LDHC* high or low expressing tumors by CTL level. **D)** Kaplan Meier survival curves of melanoma patients receiving PD-1 blockade, illustrating overall and progression-free survival in relation to tumor *LDHC* expression.

### Silencing of LDHC improves T cell activation and cytolytic function

To investigate the role of LDHC tumor expression in regulating T cell function, we first generated breast cancer cell models with varying levels of LDHC expression by knocking down *LDHC* in two basal-like cell lines (MDA-MB-468, BT-549) and one Her2-enriched cell line (HCC-1954). Downregulation of LDHC expression was confirmed at the RNA and protein level in all three cell lines **(Fig 2A-B)**. Next, we co-cultured *LDHC*-silenced tumor cells and their control counterparts with peripheral blood lymphocytes and assessed IFN-γ secretion using the ELISpot assay. Silencing of LDHC significantly increased IFN-γ production, a marker of T cell activation, across all three cell lines **(Fig 2C)**. In addition, we observed a significant increase in immune cell-mediated cancer cell killing following LDHC silencing in all cell lines **(Fig 2D)**.

**Figure 2.**
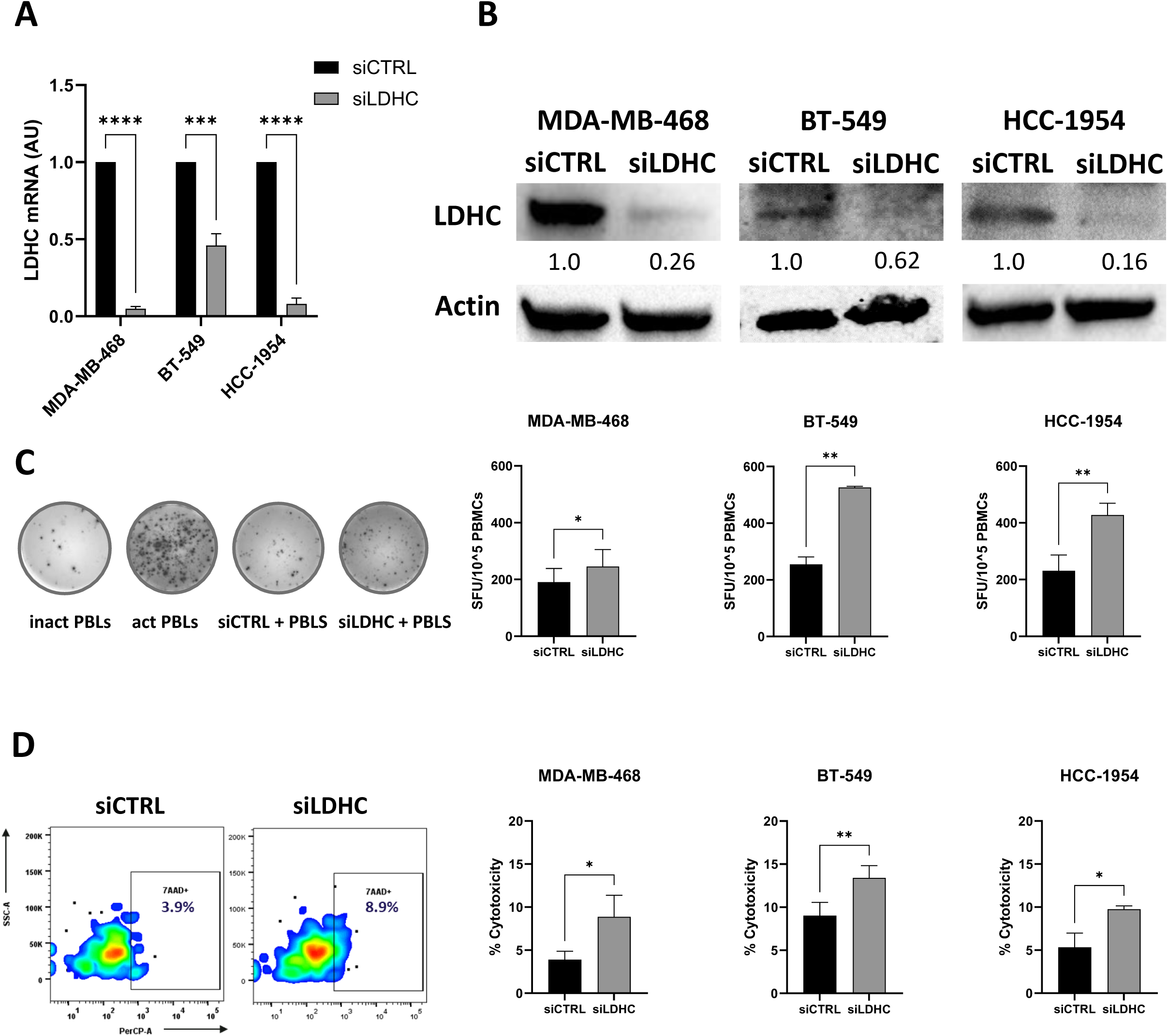
LDHC knockdown enhances T cell function and cytolytic activity. **A)** Validation of LDHC silencing efficiency in three breast cancer cell lines using real-time qRT-PCR (normalized to expression of housekeeping gene *RPLPO*). Bar charts are representative of combined data from 3 independent experiments. Statistical analysis comparing siCTRL vs siLDHC performed using unpaired Student’s t-test. **B)** Confirmation of LDHC knockdown by western blotting. Representative image from three independent experiments. **C)** Analysis of T cell activity as measured by IFN-γ ELISpot after short-term direct co-culture of cancer cells with HLA-matched peripheral blood lymphocytes. Combined data from 3 independent experiments, each performed with one donor (biological replicate), and one representative image (MDA-MB-468 cells) are depicted. **D)** Cell viability flow cytometry analysis after short-term direct co-culture. Non-viable cancer cells were gated as positive for Qtracker and 7-AAD staining. Combined data from 3 independent experiments, each performed with one donor (biological replicate), and one representative flow cytometry plot (MDA-MB-468 cells) are shown. Bar charts represent mean with standard error of mean (±SEM) from three independent replicates. *p ≤ 0.05, **p ≤ 0.01, ***p≤0.001, **** p≤0.0001

### Tumor LDHC expression alters the production of tumor-derived inflammatory mediators and expression of immune checkpoint ligands

Following our observations that decreased *LDHC* expression in breast cancer cells was associated with enhanced T cell functionality, we performed an in-depth analysis of LDHC- associated changes in immune-related molecules. For these analyses, we utilized the basal-like MDA-MB-468 breast cancer cell line which demonstrated robust LDHC knockdown efficiency and the highest increase in T cell-mediated cancer cell cytotoxicity following LDHC silencing. Using a 23-plex bead-based immunoassay, we found a significant increase in cancer cell-derived GM-CSF, IFN-γ, MCP-1, and CXCL1 levels following knockdown of LDHC, whereas the levels of IL-6 and Gal-9 were significantly reduced **(Fig 3A)**. Protein-protein interaction network analysis **(Fig 3B)** revealed that the upregulated proteins form a high-confidence network (PPI enrichment p value=2.5e-8). Gene ontology analysis indicated that these upregulated molecules are mainly involved in processes supporting an immune favorable environment, including cytokine and chemokine-mediated signaling, phagocytosis, cell proliferation, macrophage differentiation, chemotaxis and STAT signaling **(Fig 3B)**. Furthermore, the increase in IFN- γ levels is in accordance with the increased T cell activation and cytotoxic activity we observed in direct co-cultures **(Fig 2C-D)**. In turn, the reduction in secreted IL-6 and Gal-9 levels may contribute to enhanced anti-tumor immunity by alleviating the pro-tumorigenic and immunosuppressive effects of the pleiotropic cytokine IL-6 and by impeding the interaction of Gal-9 with the immune checkpoint receptor TIM3 on T cells.

**Figure 3.**
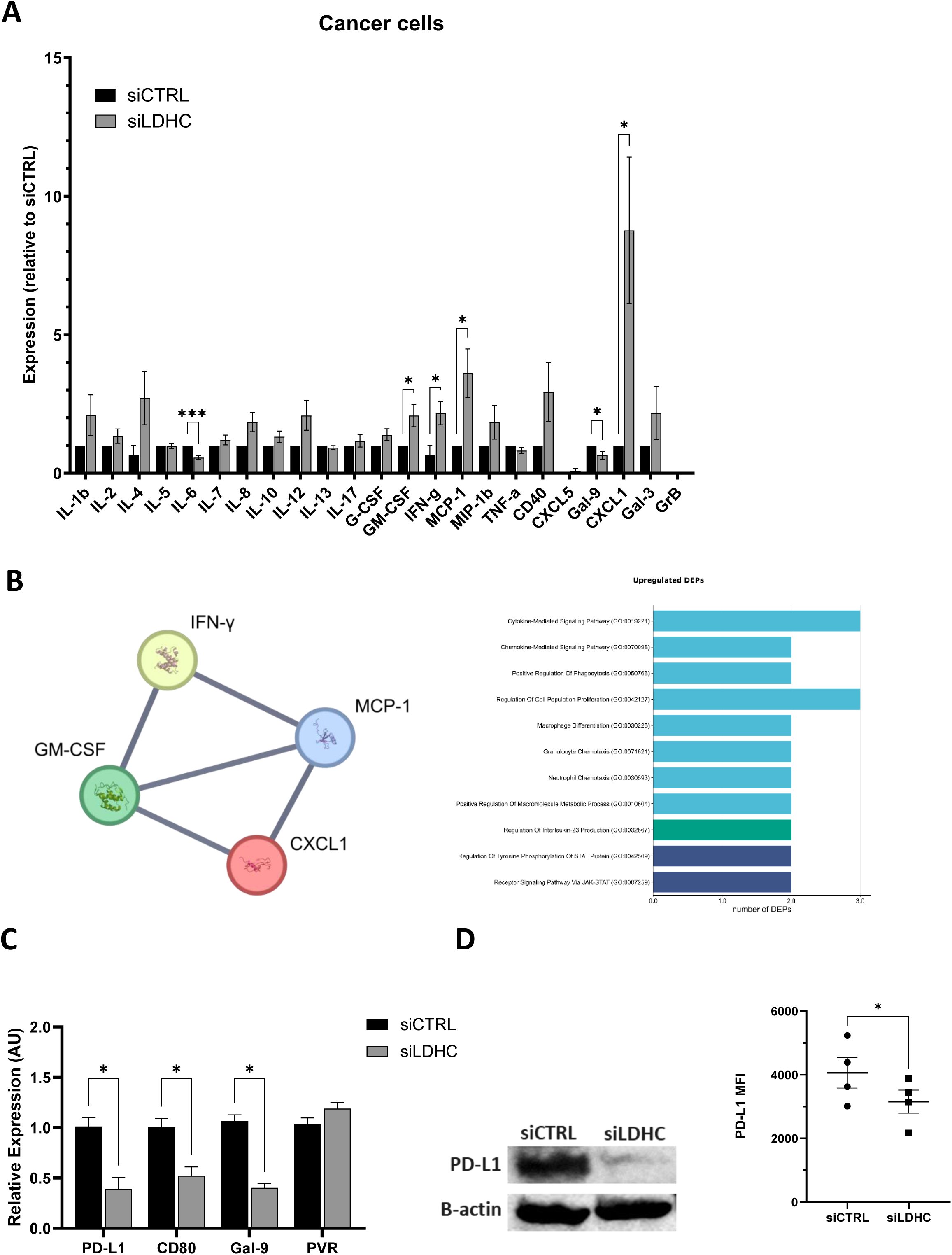
LDHC knockdown alters tumor-derived cytokine and chemokine secretion and immune checkpoint expression in MDA-MB-468 breast cancer cells. **A)** Cytokine and chemokine levels measured by 23-plex using 72-hour conditioned media from cancer cells following LDHC knockdown. Protein levels are expressed as mean fold-change relative to siCTRL. **B)** Protein-protein interaction network and gene ontology analysis of proteins with increased expression in LDHC-silenced cells. **C)** Analysis of immune checkpoint expressions using real time qRT-PCR, western blotting and flow cytometry. RNA expression data was normalized to *RPLPO* expression and plotted as mean fold-change relative to siCTRL. For western blotting, β-actin protein expression was used as a loading control. Data is representative of a minimum of three independent experiments. Bar charts represent mean with standard error of mean (±SEM). Statistical analysis performed using Student’s t-test. * p ≤ 0.05, *** p≤0.001. MFI, mean fluorescence intensity; DEP, differentially expressed protein.

In addition to soluble inflammatory molecules, stimulatory and inhibitory immune checkpoints play an important role in regulating anti-tumor immunity. Hence, we assessed the tumor cell expression of four distinct immune checkpoint ligands; *PD-L1*, *CD80*, *GAL-9,* and *PVR*, which interact with the clinically relevant receptors *PD-1*, *CTLA-4, TIM3*, and *TIGIT* on T cells. Silencing LDHC significantly downregulated the mRNA expression of *PD-L1*, *CD80* and *GAL-9* in MDA-MB-468 breast cancer cells **(Fig 3C)**. Further analysis of BT-549 and HCC- 1954 demonstrated a significant downregulation of *PD-L1* expression in both cell lines and a trend towardsreduced *GAL-9* in HCC-1954 cells **(Fig S1)**. Given the significant clinical benefit of PD-1/PD-L1 blockade in multiple cancers, we further validated the downregulation of PD- L1 in *LDHC*-silenced MDA-MB-468 cells using western blotting and flow cytometry **(Fig 3C)**.

### Immunomodulatory effects of aberrant LDHC tumor expression require direct cell-cell contact of cancer cells with immune cells

Since we found that aberrant expression of LDHC in tumor cells impacts T cell activity **(Fig 2C-D)** and dysregulates the secretion of tumor-derived pro-inflammatory molecules **(Fig 3A- B)**, we sought to assess the effects of LDHC silencing on cytokine levels in cancer cell-PBL co-culture models. Using an indirect co-culture model, we found a mere decrease in Gal-9 levels (p<0.01), similar to what we observed in cancer cell monocultures, suggesting that modulation of the anti-tumor immune response in co-culture models requires cell-cell contact **(Fig 4A).** Using direct co-culture of *LDHC*-silenced cancer cells with PBLs, we observed multiple changes including a significant decrease in the pro-tumorigenic cytokines IL-1β (p<0.01), IL-4 (p<0.01), and IL-6 (p=0.03), alongside IFN-γ (p=0.01) and MIP-1b (p=0.04) and an increase in CXCL1 (p=0.04) **(Fig 4B).** Network analysis demonstrated high-confidence interconnections (PPI enrichment p value=1.21e-12) between each of the five downregulated molecules **(Fig 4C)**.

**Figure 4.**
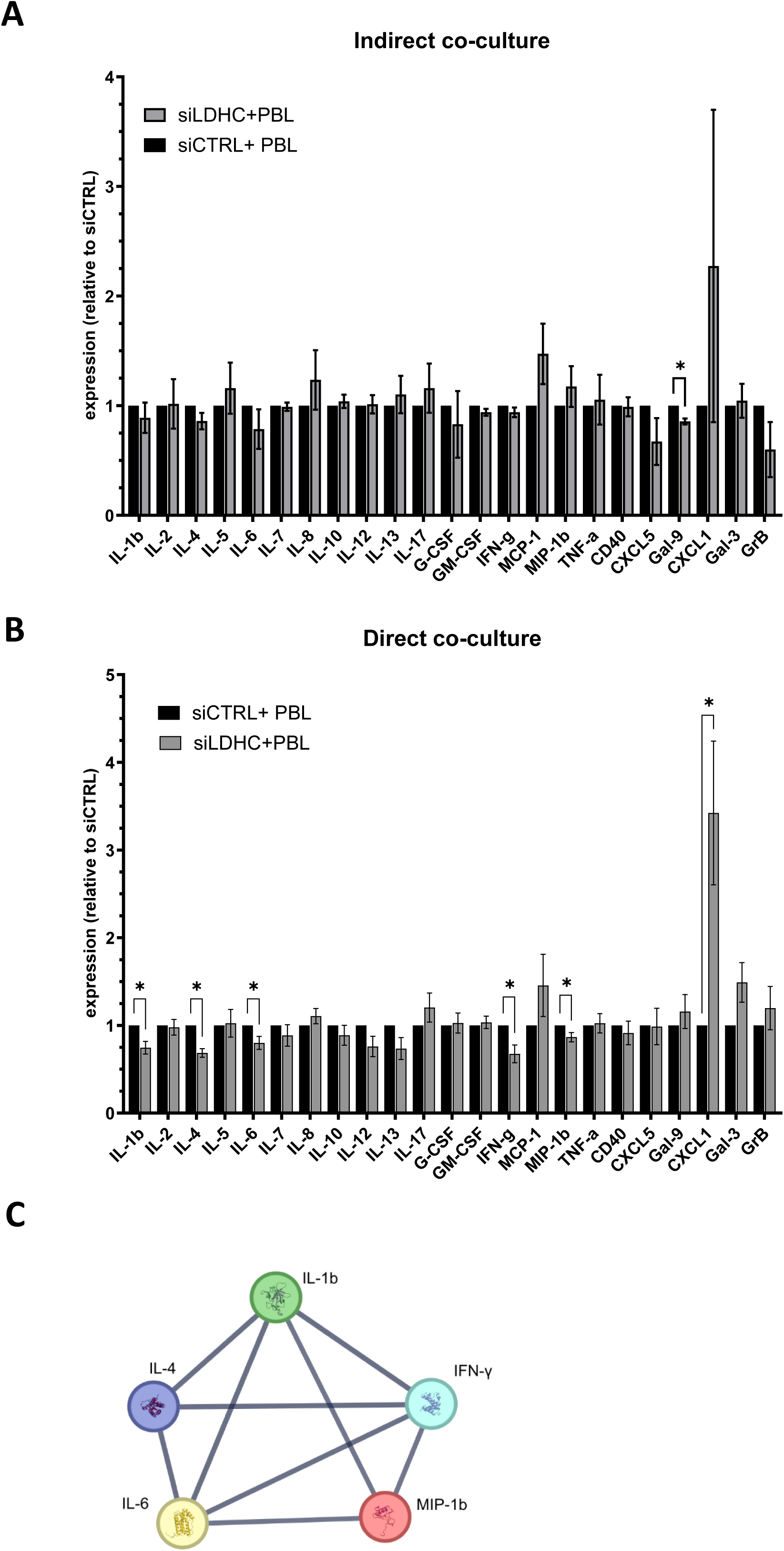
LDHC knockdown in MDA-MB-468 breast cancer cells alters soluble cytokine and chemokine levels in indirect and direct co-cultures. **A)** Cytokine and chemokine levels measured by 23-plex using 72 hours conditioned media from indirect co-cultures. **B)** Cytokine and chemokine analysis of 72-hour conditioned media from direct cocultures. **C)** Protein-protein interaction network of downregulated proteins in LDHC-silenced cells. Protein levels are expressed as mean fold-change relative to siCTRL. Combined data from a minimum of three independent experiments, each performed with one donor (biological replicate), are shown. Bars indicate mean with standard error of mean (±SEM). Statistical analysis performed using unpaired Student’s t-test. * p ≤ 0.05. DEP, differentially expressed protein.

Additionally, we assessed the expression of immune checkpoint ligands and receptors. Analysis of CD8+ T cell surface expression revealed a significant reduction in the number of cells expressing CTLA-4 in direct co-cultures and PD-1 in indirect co-cultures **(Fig 5A-B)**. Furthermore, the expression of the immune checkpoint molecules itself was altered upon co-culture. For instance, TIGIT, TIM3 and VISTA expression was downregulated by 72 hours of direct co-culture with *LDHC-*silenced cancer cells, while CTLA-4 expression was reduced in indirect co-cultures **(Fig 5A-B, Fig S2A-B)**. Looking at the cancer cells, we find that the number of *LDHC*-silenced cancer cells expressing PD-L1, PD-L2 and CD80 is reduced and that the cell surface expression of PD-L2, Gal-9, PVR, HLA-DR and VISTA is decreased as well **(Fig 6A-B, Fig S2C)**. Together, these findings suggest that silencing of LDHC in cancer cells disrupts multiple immune checkpoint signaling axes, resulting in enhanced T cell activity.

**Figure 5.**
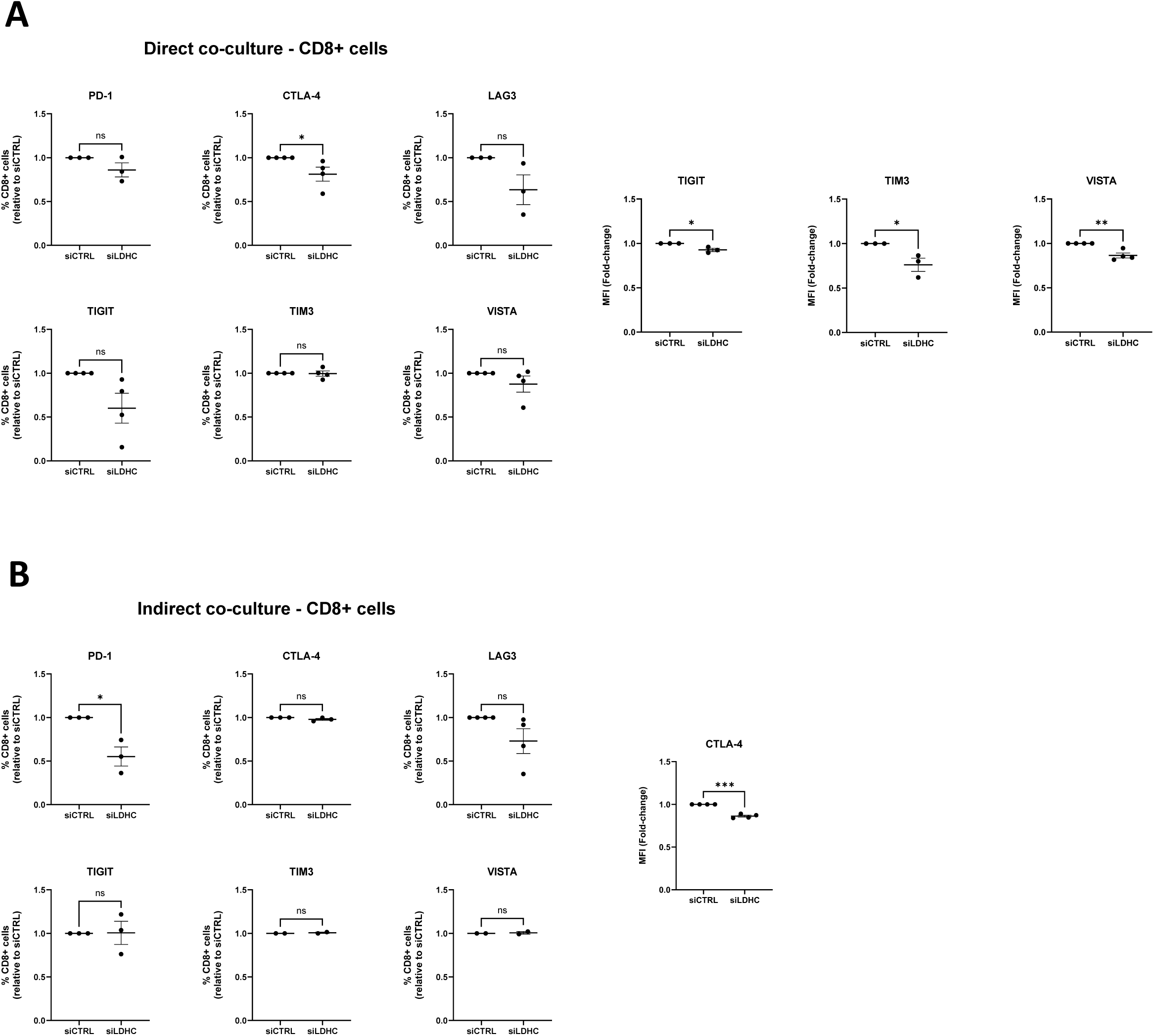
LDHC knockdown reduces expression of immune checkpoint receptors on CD8+ T cells following direct and indirect coculture with MDA-MB-468 breast cancer cells. **A)** Flow cytometry analysis to assess the frequency of CD8+ T cells expressing PD-1, CTLA-4, LAG-3, TIGIT, TIM3 and VISTA after 72 hours of direct co-culture *(left)*, and their expression of TIGIT, TIM3 and VISTA *(right)*. **B)** Flow cytometry analysis of the number of CD8+ T cells expressing immune checkpoint receptors after 72 hours of indirect co-culture *(left)* and of CTLA-4 expression *(right)*. Dot plots represent mean fold-change relative to siCTRL with standard error of mean (±SEM). Combined data from multiple independent experiments, each performed with one PBL donor (biological replicate), are shown. Statistical analysis performed using paired Student’s t-test. * p ≤ 0.05, ** p ≤ 0.01, *** p ≤ 0.001.

**Figure 6.**
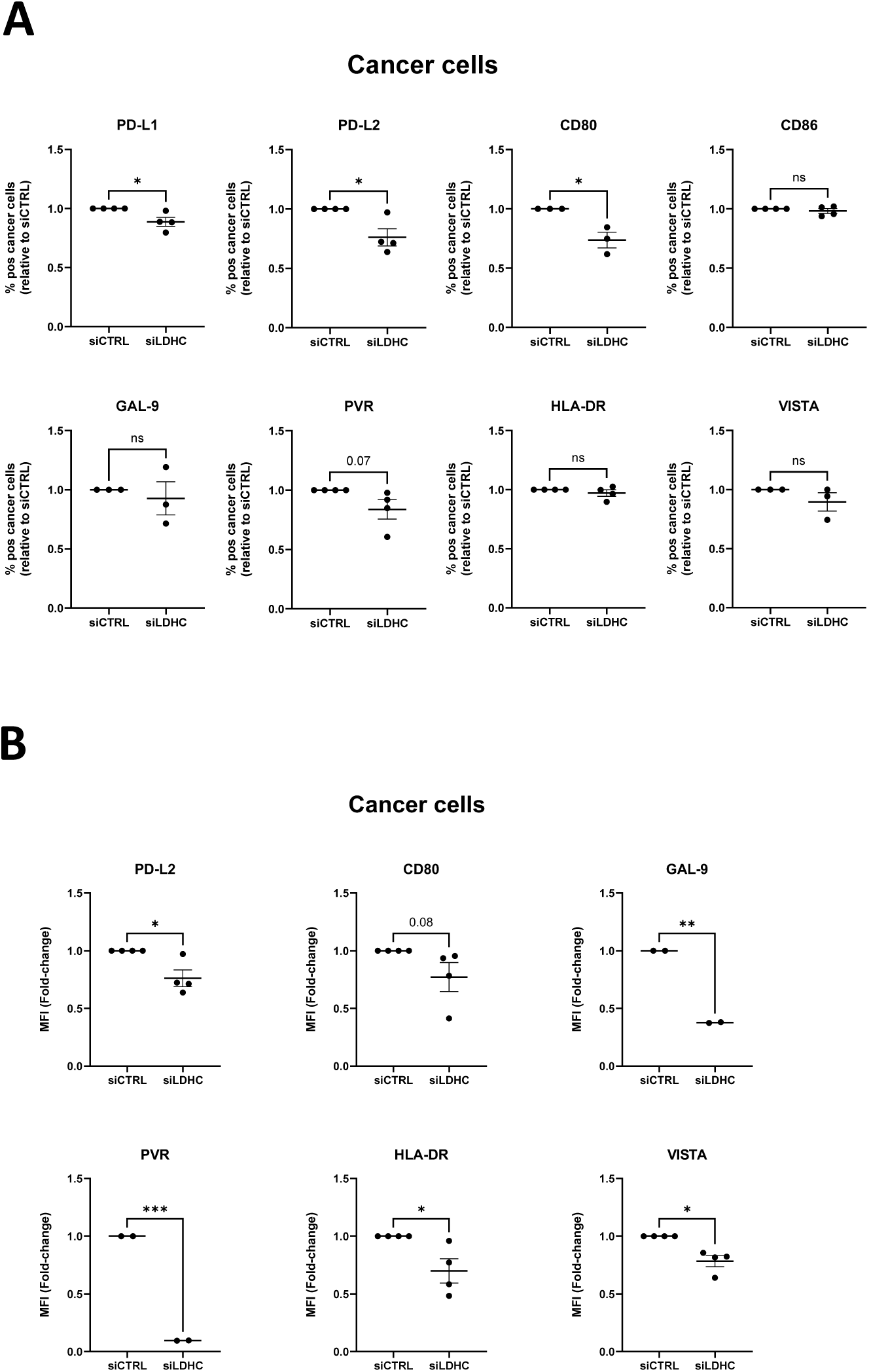
LDHC knockdown reduces expression of immune checkpoint ligands on MDA-MB-468 cancer cells following indirect co-culture. **A)** Flow cytometry analysis of the number of cancer cells expressing PD-L1, PD-L2, CD80, CD86, GAL-9, PVR, HLA-DR and VISTA after 72 hours of indirect co-culture. **B)** Flow cytometry analysis of the expression of immune checkpoint ligands on cancer cells. Dot plots represent mean fold-change relative to siCTRL with standard error of mean (±SEM). Combined data from multiple independent experiments, each performed with one PBL donor (biological replicate), are shown. Statistical analysis performed using paired Student’s t-test. * p ≤ 0.05, ** p ≤ 0.01, *** p ≤ 0.001.

## DISCUSSION

Traditionally, cancer treatment has focused on reducing or eliminating tumor burden through surgery, radiation therapy and chemotherapy. However, advances in our understanding of cancer biology and immunology have revolutionized cancer treatment, leading to the integration of targeted therapies and immunotherapy alongside traditional treatment approaches. Immunotherapy with immune checkpoint inhibitors targeting PD-1/PD-L1 and CTLA-4 in particular has greatly improved clinical outcomes of patients with melanoma, non-small cell lung cancer, and renal cell carcinoma [31–33]. Despite these advances, the success of immunotherapy is limited to a subset of patients, with a significant proportion of patients either failing to respond or developing severe immune-related adverse events [34]. To enhance the efficacy of immunotherapy, it is crucial to gain insights into the intricate interactions between tumor cells and their microenvironment, including stromal cells and immune cells.

In the present study, we investigated how targeting LDHC in cancer cells might impact anti-tumor immunity, potentially offering a novel dual approach to target tumor cells and promote the immune response. We specifically focused on basal-like and Her2-enriched breast tumors as we previously found that they exhibit a higher expression of *LDHC* compared to luminal breast tumors [19]. Here, we show through TIDE analysis that increased *LDHC* expression in these tumors is associated with T cell dysfunction. In accordance, we found that silencing of LDHC in three distinct breast cancer cell lines enhanced T cell activation and cytolytic activity. Comprehensive analysis of 23 cytokines and chemokines revealed an increase in tumor-derived GM-CSF, IFN-γ, MCP-1 and CXCL1, alongside a decrease in IL-6 and Gal-9, which supports the enhanced T cell activity we observed following LDHC knockdown. Furthermore, we found a reduction in PD-L1 expression on the cell surface of *LDHC*-silenced tumor cells, likely impairing PD-L1/PD-1 signaling and resulting in improved T cell activation.

Analysis of cancer cell-immune cell co-cultures revealed how tumor LDHC expression differentially affects the expression of soluble inflammatory mediators through direct cell-cell contact and indirect interactions. We observed that direct co-culture alters the expression levels of several cytokines following LDHC knockdown, while only one analyte was affected in indirect co-cultures, suggesting that tumor cell LDHC expression requires direct cell-cell contact to significantly influence immune cell activity in co-culture conditions. Further, we found that direct co-culture of LDHC-silenced breast cancer cells with peripheral blood lymphocytes induces early T cell activation, as shown by IFN-γ ELISpot and 7-AAD cytotoxicity assays after 4 hours of co-culture. Further, we show that this early immune response is followed by a decrease in IL-1β, IL-4 and IL-6 and an increase in CXCL1 at 72 hours co-culture. Together, IL-1β, IL-4 and IL-6 are well-known for their tumor promoting effects, which include enhancing tumor cell proliferation and survival, supporting cancer cell stemness, and facilitating invasion and angiogenesis through autocrine signaling or crosstalk with tumor infiltrating immune cells [35–37]. On the other hand, elevated expression of CXCL1 contributes to improved tumor control through the recruitment and activation of neutrophils [38]. This suggests that targeting tumor LDHC expression could provide a novel strategy to promote early T cell activation, disrupt tumor-promoting cytokines and enhance the activity of tumor-suppressive cytokines. Thus, further studies are needed to better comprehend the effects of targeting LDHC on the initiation and maintenance of anti-tumor immune responses. In addition, co-culture models involving different immune cell subpopulations will be crucial to dissect the interactions between LDHC expressing tumor cells and different components of the immune response. Of note, the significant reduction in soluble Gal-9 levels in cancer cell monocultures and indirect co-cultures suggests that secretion of Gal-9 is primarily regulated by paracrine signaling rather than by direct interactions with immune cells. As such, silencing LDHC and the subsequent decrease in soluble Gal-9 may result in reduced binding of Gal-9 to the inhibitory immune checkpoint receptor TIM3, leading to enhanced T cell activation [39]. Furthermore, we found that LDHC knockdown results in a decrease in the expression of several immune checkpoint ligands and receptors, as well as a reduction in the number of PD-1 and CTLA-4 positive CD8+ T cells following co-culture. Together, these findings indicate that aberrant LDHC tumor expression is associated with immune dysfunction through impairment of T cell activation and modulation of immune checkpoint signaling.

## CONCLUSIONS

Immunotherapy has dramatically changed cancer care; however, its effectiveness remains limited due to various tumor-intrinsic and microenvironmental factors. Cancer cells produce various pro-tumorigenic and immunoFigive molecules that drive tumor progression and hinder immune-mediated tumor elimination. Therefore, to improve therapeutic outcomes, a better understanding of the dynamic interactions between cancer cells and immune cells is essential to gain insight into tumor cell surface markers and tumor-derived soluble factors that influence anti-tumor immunity. Our findings suggest that targeting tumor LDHC expression could help foster a more favorable immune microenvironment, potentially enhancing responses to immunotherapy. Further studies are needed to elucidate the molecular mechanisms by which LDHC modulates immune responses, including its metabolic role in promoting immunosuppression.

## LIST OF ABBREVIATIONS

7-AAD: 7-Aminoactinomycin D
Akt: Ak strain transforming
ATCC: American Tissue Culture Collection
BRCA: breast cancer
CREB: cAMP-response element binding protein
CTA: cancer testis antigen
CTL: cytotoxic T lymphocytes
CTLA-4: Cytotoxic T-lymphocyte associated protein 4
CXCL1: chemokine (C-X-C motif) ligand 1
DEP: differentially expressed protein
ELISPot: Enzyme-linked immunosorbent spot
EPIC: Estimating the Proportions of Immune and Cancer cells
FBS: fetal bovine serum
FDA: Food and drug administration
Gal-9: galectin 9
GM-CSF: Granulocyte-macrophage colony-stimulating factor
GO: gene ontology
GSK-3B: Glycogen Synthase Kinase 3 Beta
HLA: human leukocyte antigen
IFN-γ: interferon gamma
IL: interleukin
IPA: ingenuity Pathway analysis
LAG-3: Lymphocyte activation gene 3 protein
LDHA: Lactate dehydrogenase A
LDHB: Lactate Dehydrogenase B
LDHC: Lactate Dehydrogenase C
LUMA: luminal A
MAGE-C2: Melanoma-associated antigen C2
MCP-1: Monocyte Chemoattractant Protein-1
MFI: mean fluorescence intensity
NK: Natural Killer
OS: overall survival
PBK: PDZ Binding Kinase
PBL: peripheral blood lymphocytes
PBMC: peripheral blood mononuclear cells
PCR: polymerase chain reaction
PD-1: Programmed cell death protein 1
PD-L1: Programmed death ligand 1
PFS: progression free survival
PI3K: Phosphoinositide 3-kinase
PPI: Protein-protein interaction
PRAME: Preferentially expressed antigen in melanoma
PVR: poliovirus receptor
RNA: Ribonucleic acid
RPLPO: ribosomal protein lateral stalk subunit P0
RPMI: Roswell Park Memorial Institute 1640
SFU: spot forming units
siCTRL or siLDHC: small interfering CTRL or LDHC RNA
Sp1: specificity protein 1
SSX2: Synovial sarcoma X breakpoint 2
STAT: Signal Transducer and Activator of Transcription
T:E: Target:Effector ratio
TCGA: The Cancer Genome Atlas
TIDE: Tumor Immune Dysfunction and Exclusion
TIGIT: T cell immunoreceptor with Ig and ITIM domains
TIM3: T cell immunoglobulin mucin-3
TIMER: Tumor IMmune Estimation Resource
TPM: transcripts per million
VISTA: V-domain immunoglobulin suppressor of T cell activation

## DECLARATIONS

### ETHICS APPROVAL AND CONSENT TO PARTICIPATE

The PBL collection protocol was approved by the Qatar Biomedical Research Institute (study approval number 2016-002, initial approval date 31st May 2017) and the Hamad Medical Corporation (study approval number 17132/17, initial approval date 5th November 2017) institutional review boards and was performed in accordance with the 1964 Helsinki Declaration and its later amendments. The study protocol was granted IRB waiver of informed consent under the condition of anonymization and no additional intervention to the participants.

### CONSENT FOR PUBLICATION

Not applicable.

### AVAILABILITY OF DATA AND MATERIALS

All data that support the findings of this study are included in this published article and its supplementary information files. Additional data can be provided upon request from the corresponding author.

### AUTHOR CONTRIBUTIONS

AN performed experiments, analyzed the data and wrote the first draft. RT performed experiments and revised the manuscript. AAK performed PD-L1 qRT-PCR experiments and revised the manuscript. HQ analyzed data and revised the manuscript. JD conceptualized the study, was responsible for funding acquisition and project administration, and revised the manuscript. All authors read and approved the final manuscript.

## ACKNOWLEDGEMENTS

Not applicable.

