## Supplementary figures and images for "Immunomodulatory effects of tumor Lactate Dehydrogenase C (LDHC) in breast cancer"

### Fig S1

#### Supplementary Figure 1

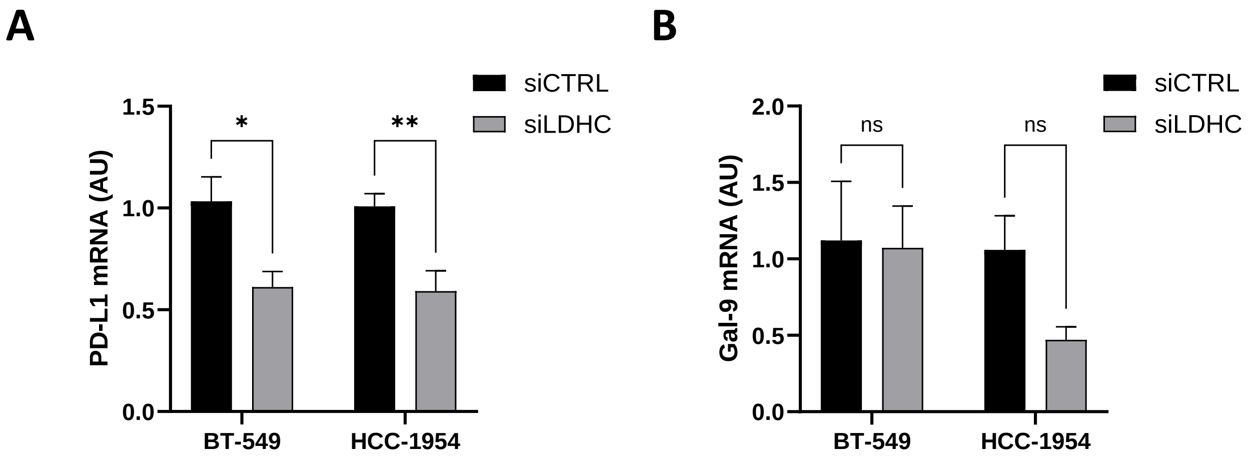

### Fig S2

#### Supplementary Figure 2

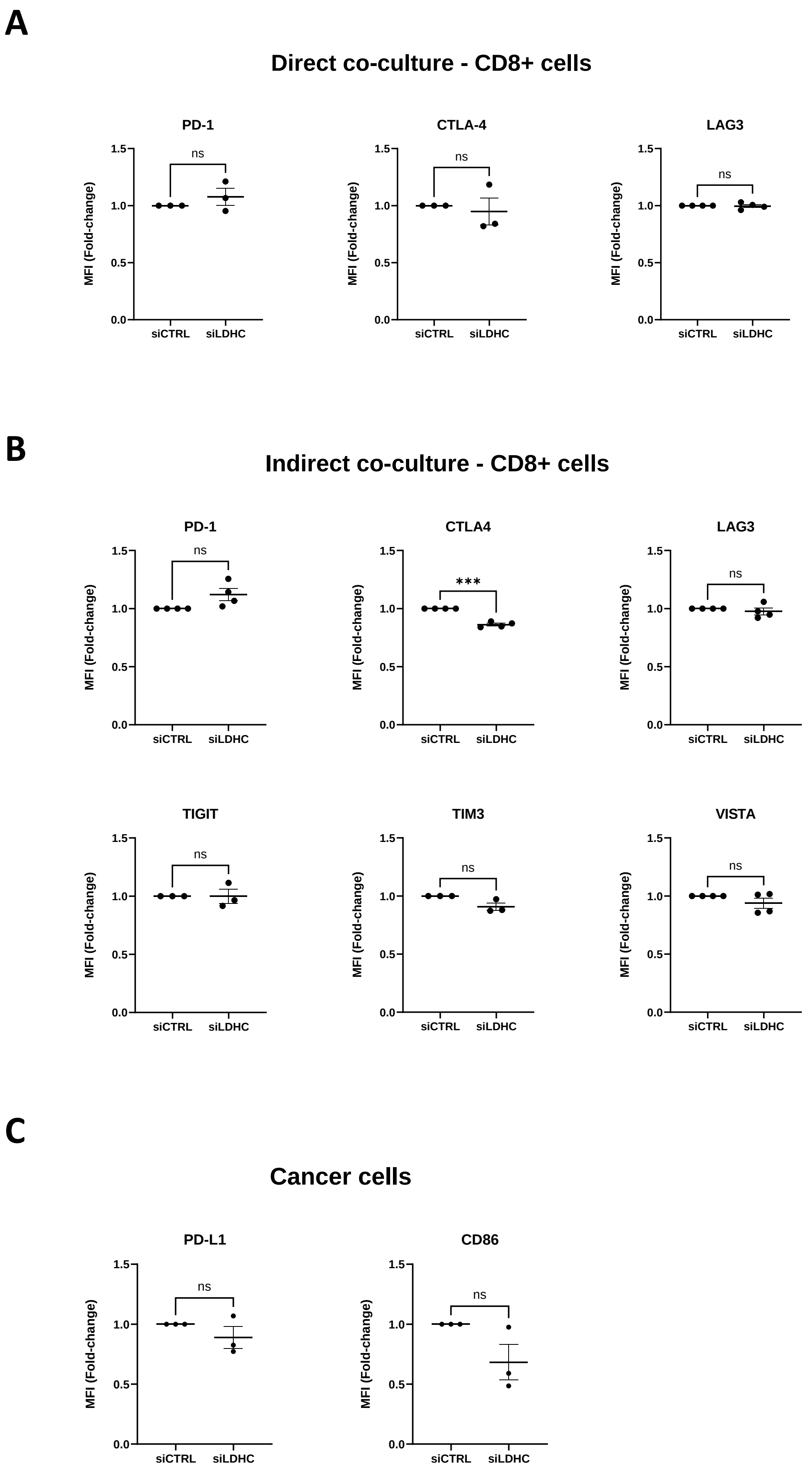
